# Activation of intracellular transport by relieving KIF1C autoinhibition

**DOI:** 10.1101/488049

**Authors:** Nida Siddiqui, Alice Bachmann, Alexander James Zwetsloot, Hamdi Hussain, Daniel Roth, Irina Kaverina, Anne Straube

## Abstract

The kinesin-3 KIF1C is a fast organelle transporter implicated in the transport of dense core vesicles in neurons and the delivery of integrins to cell adhesions. Here we report the mechanisms of autoinhibition and release that control the activity of KIF1C. We show that the microtubule binding surface of KIF1C motor domain interacts with its stalk and that these autoinhibitory interactions are released upon binding of protein tyrosine phosphatase PTPN21. The FERM domain of PTPN21 stimulates dense core vesicle transport in primary hippocampal neurons and rescues integrin trafficking in KIF1C-depleted cells. In vitro, human full-length KIF1C is a processive, plus-end directed motor. Its landing rate onto microtubules increases in the presence of either PTPN21 FERM domain or the cargo adapter Hook3 that binds the same region of KIF1C tail. This autoinhibition release mechanism allows cargo-activated transport and might enable motors to participate in bidirectional cargo transport without undertaking a tug-of-war.

## INTRODUCTION

Intracellular transport is essential for cell polarity and function. Long-distance transport of cellular cargo is mediated by microtubule-based motors, dynein and kinesin. While dynein is the main transporter towards the minus end of microtubules, most kinesins walk towards the microtubule plus end (Hancock, 2014). However, many cargoes carry motors of both directionality in order to allow directional switching when they encounter a roadblock and the relative activity of the opposite directionality motors determines the net progress of the cargo towards the cell periphery (where usually most plus ends are located) or towards the cell centre (where microtubule minus ends are abundant) (Ashkin et al., 1990, Hancock, 2014). We have previously identified the kinesin-3 KIF1C as the motor responsible for the transport of α5β1-integrins. The delivery of integrins into cellular processes such as the tail of migrating cells allows the maturation of focal adhesion sites (Theisen et al., 2012). KIF1C is also required for the formation and microtubule-induced turnover of podosomes – protrusive actin-based adhesion structures – in both macrophages and vascular smooth muscle cells (Kopp et al., 2006, Efimova et al., 2014, Bhuwania et al., 2014). KIF1C also contributes to MHC class II antigen presentation, Golgi organization and transport of Rab6-positive secretory vesicles (Lee et al., 2015, del Rio et al., 2012). In neurons, KIF1C transports dense core vesicles both into axons and dendrites and appears to be the fastest human cargo transporter (Schlager et al., 2010, Lipka et al., 2016). Consistently, human patients with missense mutations in KIF1C resulting in the absence of the protein suffer from spastic paraplegia and cerebellar dysfunction (Dor et al., 2014). Mutations that result in reduced motor function also cause a form of hereditary spastic paraplegia (Caballero Oteyza et al., 2014).

KIF1C-dependent cargo moves bidirectional even in highly polarised microtubule networks and depletion of KIF1C results in the reduction of transport in both directions (Theisen et al., 2012), suggesting that KIF1C cooperates with dynein as had been suggested for other kinesin-3 motors (Bielska et al., 2014). To begin to unravel how KIF1C contributes to bidirectional cargo transport, we need to identify the mechanisms that switch KIF1C transport on and off. Most kinesin-3 motors are thought to be activated by a monomer-dimer switch, whereby cargo binding releases inhibitory intramolecular interactions of neck and stalk regions by facilitating the dimerisation of the neck coil and other coiled-coils regions in the tail (Al-Bassam et al., 2003, Soppina et al., 2014, Rashid et al., 2005, Tomishige et al., 2002). The motor thereby transitions from an inactive, diffusive monomer to a processive dimer. In the alternative tail block model, the motors are stable dimers, but regions of the tail interact with the motor or neck domains and interfere with motor activity until cargo binding occupies the tail region and releases the motor (Yamada et al., 2007, Farkhondeh et al., 2015, Yoshimura et al., 2010).

Here we show that KIF1C is a stable dimer that is autoinhibited by intramolecular interactions of the stalk domain with the microtubule binding interface of the motor domain. We demonstrate that protein tyrosine phosphatase N21 (PTPN21) activates KIF1C by binding to the tail region. This function does not require catalytic activity of the phosphatase and its N-terminal FERM domain alone is sufficient to stimulate the transport of KIF1C cargoes in cells as well as increasing the landing rate of KIF1C on microtubules in vitro. The cargo adapter Hook3 binds KIF1C in the same region and activates KIF1C in a similar way, suggesting that both cargo binding and regulatory proteins might contribute to the directional switching of KIF1C-dynein transport complexes.

## RESULTS

### Kif1C is an autoinhibited dimer

To determine the mechanism of KIF1C regulation, we first aimed to determine its mode of autoinhibition. The monomer-dimer-switch and tail-block models can be distinguished by determining the oligomeric state of the motor. Thus we purified recombinant full-length human KIF1C from insect cells and performed classic hydrodynamic analysis using glycerol gradients and size exclusion chromatography (Figure 1a-d). The sedimentation coefficient and Stokes radius were determined in comparison to standard proteins at different salt concentrations. We find that the apparent molecular weight determined both at physiological levels of salt (150 mM) and in high (500 mM) salt buffer, is consistent with KIF1C being a dimer (apparent MW = 251 ± 56 kD, expected dimer MW = 308 kD). Consistent with this, the majority of KIF1C molecules bound to microtubules in the presence of non-hydrolysable AMPPNP show two bleach-steps and the distribution of bleach steps found fits best to a simulation of 89% dimers and 11% tetramers (Figure 1e-f), assuming that about 80% of GFPs are active, which is realistic based on previous findings (Ulbrich and Isacoff, 2007). Interestingly, KIF1C elongates at increasing salt concentrations from a moderately elongated conformation (frictional coefficient of 1.5) at physiological salt to highly elongated (frictional coefficient = 2.0) at 500 mM (Fig. 1d), suggesting that intramolecular electrostatic interactions might hold the KIF1C dimer in a folded, auto-inhibited state. To determine the interaction surfaces involved in a possible auto-inhibited state, we performed crosslink mass spectrometry. Purified KIF1C was treated with the 11Å crosslinker BS3 or with the zero length crosslinker EDC, digested with trypsin and then subjected to tandem mass spectrometry analysis. Crosslinked peptides were identified using StavroX (Gotze et al., 2015). Only crosslinked peptides whose identity could be verified by extensive fragmentation with none or few unexplained major peaks were retained (Figure S1). These high-confidence crosslinks were between K273-K591, K273-K645 and K464-K645, K464-E606 and E644-K854 (Figures 1g, Figure S1), showing that the end of the FHA domain and the third coiled-coil contact the motor domain near K273. This residue is within alpha-helix 4 at the centre of the microtubule interaction interface of the motor domain (Scarabelli et al., 2015) (Figure 1h). Blocking this interface with an intramolecular tail interaction therefore prevents microtubule binding and provides a plausible mechanism for auto-inhibition.

**Figure 1:**
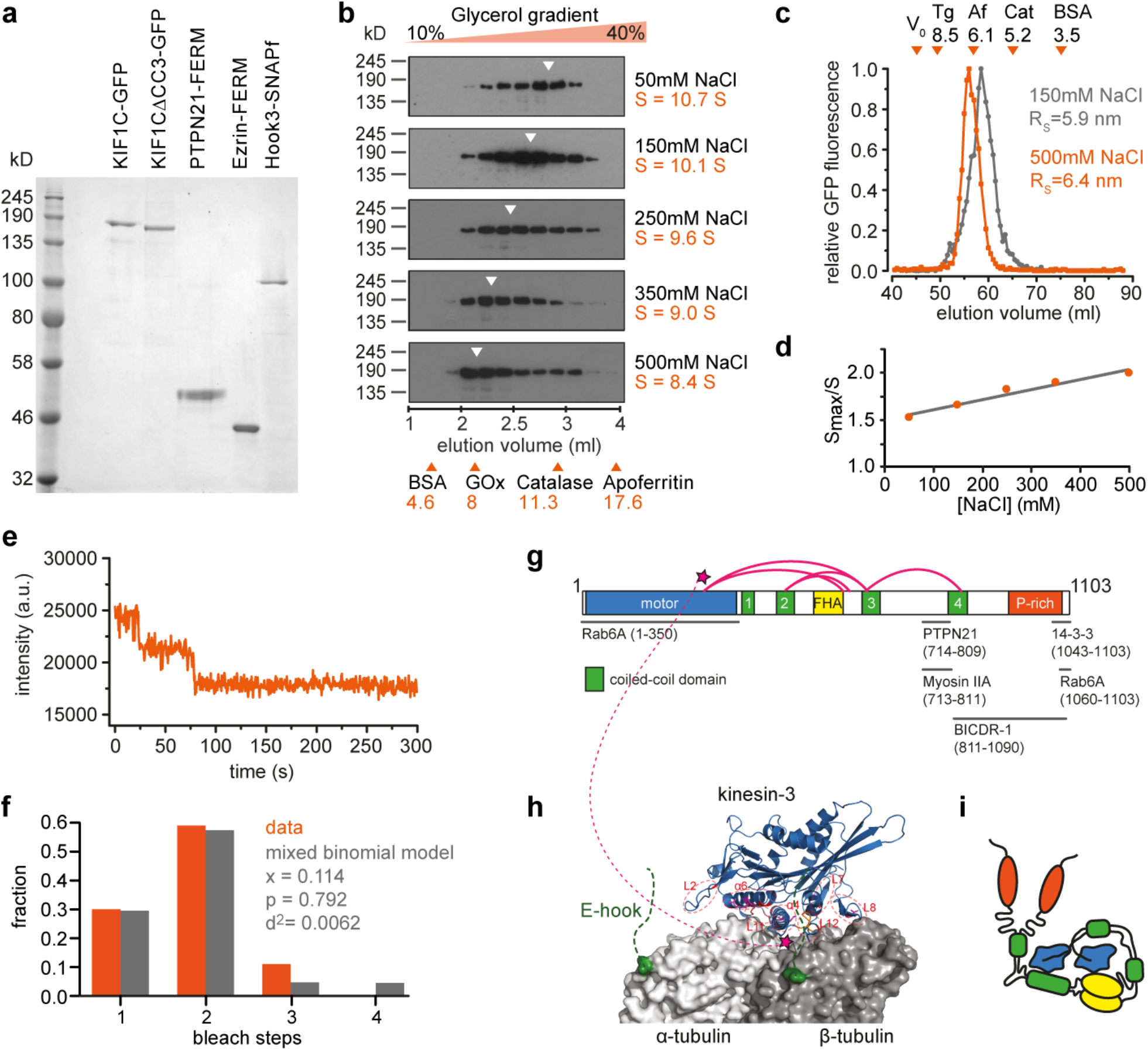
KIF1C is an autoinhibited dimer. **(a)** Coomassie stained SDS-PAGE gel of purified proteins used in this study. **(b)** Fractions from glycerol gradients of KIF1C-GFP at different salt concentrations as indicated. Elution peaks of standard proteins are indicated with orange arrowheads. GOx, glucose oxidase; BSA, bovine serum albumin. **(c)** Size exclusion chromatography of KIF1C-GFP at 150 mM NaCl (grey) and 500 mM NaCl (orange). Elution peaks of standard proteins (Tg, thyroglobulin; Af, apoferritin; Cat, catalase; BSA, bovine serum albumin) and void volume V_0_ are indicated by orange arrowheads. **(d)** Frictional coefficient of KIF1C-GFP at different salt concentrations indicating that KIF1C elongates with increasing ionic strength. **(e-f)** Bleach curve of KIF1C-GFP on microtubules showing discrete steps in fluorescent decay in e. Experimentally determined bleach steps are shown in f together with best fit to a mixed binomial model of dimers and tetramers with x being the fraction of tetramer and p the fraction of active GFP molecules. n=108 motors. **(g)** Schematic primary structure of KIF1C with motor domain (blue), coiled-coil domains (green), FHA domain (yellow) and Proline-rich domain (orange). Crosslinks identified using mass spectrometry after treatment with BS3 are shown as magenta loops. See supplementary figure S1 for ion fragmentation spectra. Published binding sites for KIF1C interactors are indicated below. **(h)** Motor domain of related KIF1A on tubulin. The region in the motor domain that interacts with KIF1C stalk is indicated by magenta stars. **(i)** Hypothetical model of autoinhibited KIF1C conformation based on identified crosslinks.

### PTPN21 stimulates transport of KIF1C cargoes

As our biochemical analysis provided strong evidence for a tail-blocked dimer configuration of KIF1C (Figure 1i), we next aimed to identify a regulator that could activate the motor. We reasoned that such a regulator would be required for KIF1C function in cells and thus would phenocopy KIF1C depletion. As readout we used the reduced podosome number in vascular smooth muscle cells that we reported previously (Efimova et al., 2014) and first tested the two proteins identified to interact with the KIF1C stalk domain in the proximity of the identified crosslinks, non-muscle Myosin IIA (Kopp et al., 2006) and protein tyrosine phosphatase N21 (PTPN21, also known as PTPD1) (Dorner et al., 1998) (Figure 1g). Myosin IIA inhibition did not phenocopy KIF1C depletion. Instead, inhibition of Myosin IIA with Blebbistatin or indirectly via inhibition of Rho kinase using Y27632 did not result in reduced podosome number, while remaining actin stress fibres were efficiently removed (Figure 2a-d). As depletion of Myosin IIA did also have opposite effects on the stability of cell tails and directional persistence of cell migration we previously reported for KIF1C depletion (Theisen et al., 2012), we excluded Myosin IIA as being involved in KIF1C activation. In contrast, PTPN21 depletion using siRNA resulted in a dramatic reduction of podosome number (Figure 2e-h). To confirm specificity of the RNAi we rescued the phenotype with HA-tagged PTPN21. Interestingly, the catalytically inactive mutant PTPN21_C1108S_ (Cardone et al., 2004) could also fully rescue the PTPN21 depletion phenotype (Figure 2e-f), suggesting that a scaffold function is required rather than phosphatase activity.

**Figure 2:**
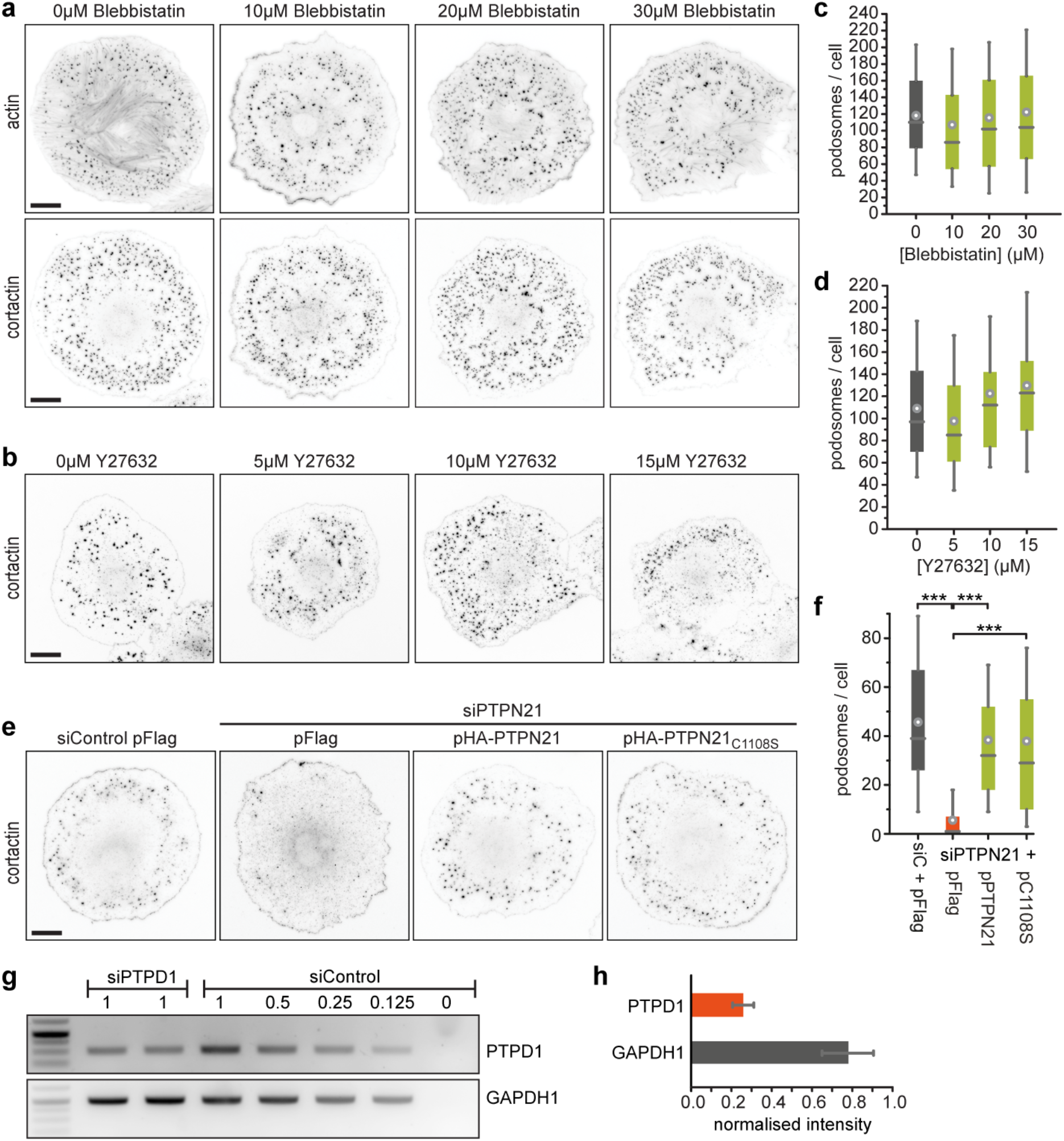
PTPN21, but not MyosinIIA is required for podosome formation. **(a-b)** A7r5 cells treated with 5 μM PDBu and different concentrations of Blebbistatin or Y27632 for 1 hour before staining for actin and cortactin. Scale bars 20 μm. **(e)** A7r5 cells transfected with siControl and siPTPN21 and either control plasmid (pFlag) or an RNAi-protected construct of wildtype PTPN21 or a catalytically inactive mutant PTPN21_c1108s_ were treated with 5 μM PDBu for 1 hour. Scale bar 20 μm. **(c,d,f)** Quantification of podosome numbers under different experimental conditions as indicated. n = 90 cells pooled from 3 independent experiments. Bar graphs show quartiles with 10/90% whiskers. Statistical significance with p>0.05 is indicated with asterisks, *** represents p<0.0005 (Mann-Whitney U-test). **(g-h)** RT-PCR from random-primed cDNA of A7r5 cells treated with siRNA as indicated. For each experiment, duplicates of siPTPN21 and 5 different concentrations of cDNA from siControl were analysed. Quantification of RT-PCR band intensities for siPTPN21 relative to siControl standard curve from 4 independent experiments is shown as mean ± SEM.

Strikingly, expression of PTPN21_C1108S_ also efficiently compensated for partial depletion of KIF1C and fully rescued podosome formation (Figure 3a-c). This suggests that PTPN21 scaffolding could activate the remaining pool of KIF1C, which is reduced to about 25-30% in cells treated with siKIF1C-2 compared to control cells (Efimova et al., 2014, Theisen et al., 2012). An N-terminal 378 amino acid fragment of PTPN21, which contains a FERM domain, was previously shown to be sufficient to interact with KIF1C (Dorner et al., 1998). Therefore we tested this construct, and found that the PTPN21 FERM domain alone was sufficient to rescue the KIF1C depletion phenotype (Figure 3a-c). To explore this further, we investigated more directly whether PTPN21 affects KIF1C-dependent transport processes. We have shown previously that KIF1C is required for the bidirectional transport of integrin-containing vesicles in migrating RPE1 cells (Theisen et al., 2012). The depletion of KIF1C results in a reduction of both plus-end and minus-end directed transport and an increase in stationary vesicles. This can be fully rescued by expression of either wildtype PTPN21 or the catalytically inactive PTPN21_C1108S_ mutant. PTPN21_FERM_ was not only able to reactivate integrin vesicle transport in KIF1C-depleted cells, but also increased motility beyond those found in control cells (Figure 3d-e). To test whether the ability of PTPN21_FERM_ to activate KIF1C-dependent transport universally applies, we next isolated primary hippocampal neurons and observed the transport of dense-core vesicles labelled with NPY-RFP in the presence of either a control plasmid or pFERM. Consistent with a function to activate KIF1C-dependent transport, we observed a dramatically increased frequency of vesicles passing through the neurites in cells expressing the FERM domain of PTPN21 (Figure 3f-g).

**Figure 3:**
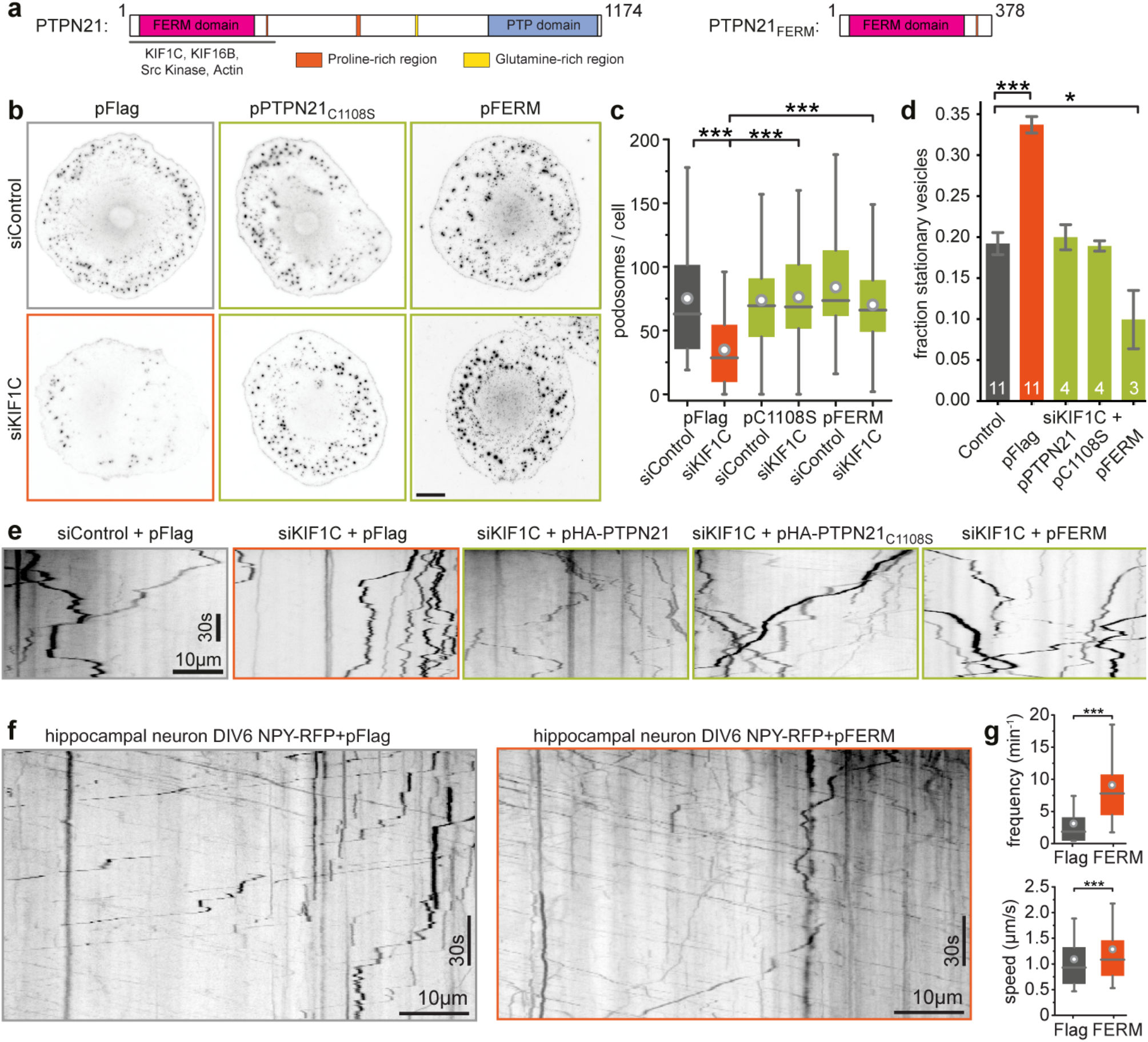
PTPN21 activates intracellular transport. **(a)** Primary structure of PTPN21 and N-terminal fragment used in this study. The region identified to interact with kinesin, actin and Src kinase is indicated below. **(b-c)** A7r5 cells treated with 5μM PDBu for 1 hour and stained for cortactin as a marker for podosomes. Expression of catalytically inactive PTPN21 or just the FERM domain rescues the podosome formation phenotype of KIF1C-depleted cells. n=60 cells. Kolmogorov-Smirnov: *** p<0.0001. Scale bar 20μm. **(d-e)** Kymographs of α5-integrin vesicles in the tail of migrating RPE1 cells. KIF1C depletion results in increase of stationary vesicles (total movement of <1.5μm) that is rescued by various PTPN21 constructs. Data show mean±SEM from 3-11 independent experiments as indicated at column bottom. *** p<0.0005, * p<0.05 (t-test with Holm Šídák correction) **(f-g)** Representative kymographs show primary hippocampal neurons isolated from a P2 mouse, transfected with NPY-RFP and either pFlag (as control) or pFERM. Number of NPY-positive vesicles passing a location per minute and the average speed of vesicles is shown. Data pooled from three independent experiments. n=36, 60 neurites / 291, 1433 vesicles. *** p<0.0001 (Kolmogorov-Smirnov).

### PTPN21 relieves KIF1C autoinhibition

Our cell biological experiments identified PTPN21 as a potential activator of KIF1C. To confirm that the PTPN21 FERM domain can directly activate KIF1C and to interrogate the mechanism of activation, we reconstituted KIF1C motility *in vitro*. Hilyte647- and biotin-labelled microtubules were assembled in the presence of GTP, then stabilised with Taxol and attached to a PLL-PEG-biotin-coated coverslip. 600 pM KIF1C was added in the presence of 1mM ATP and an ATP regenerating system. At 25 °C, individual KIF1C-GFP dimers moved processively towards the plus end of the microtubule where they accumulated (Figure 4a-b). The average speed and runlength were 0.45μm/s and 8.6μm respectively (Figure 4c). We then purified recombinant PTPN21_FERM_-6His and for comparison Ezrin_FERM_-6His from *E. coli* (Fig. 1a). Consistent with being an activator, the landing rate of KIF1C motors increased by about 40% in the presence of PTPN21_FERM_ (Figure 4b-c). The frequency of observing running motors was also increased by 40%. Ezrin_FERM_, acting as negative control, did not significantly affect the landing rate or the frequency of running motors (Figure 4b-c). These findings are consistent with the idea that PTPN21 opens the KIF1C motor by binding to its tail domain and thereby relieves auto-inhibition and increases the binding rate of the motor.

**Figure 4:**
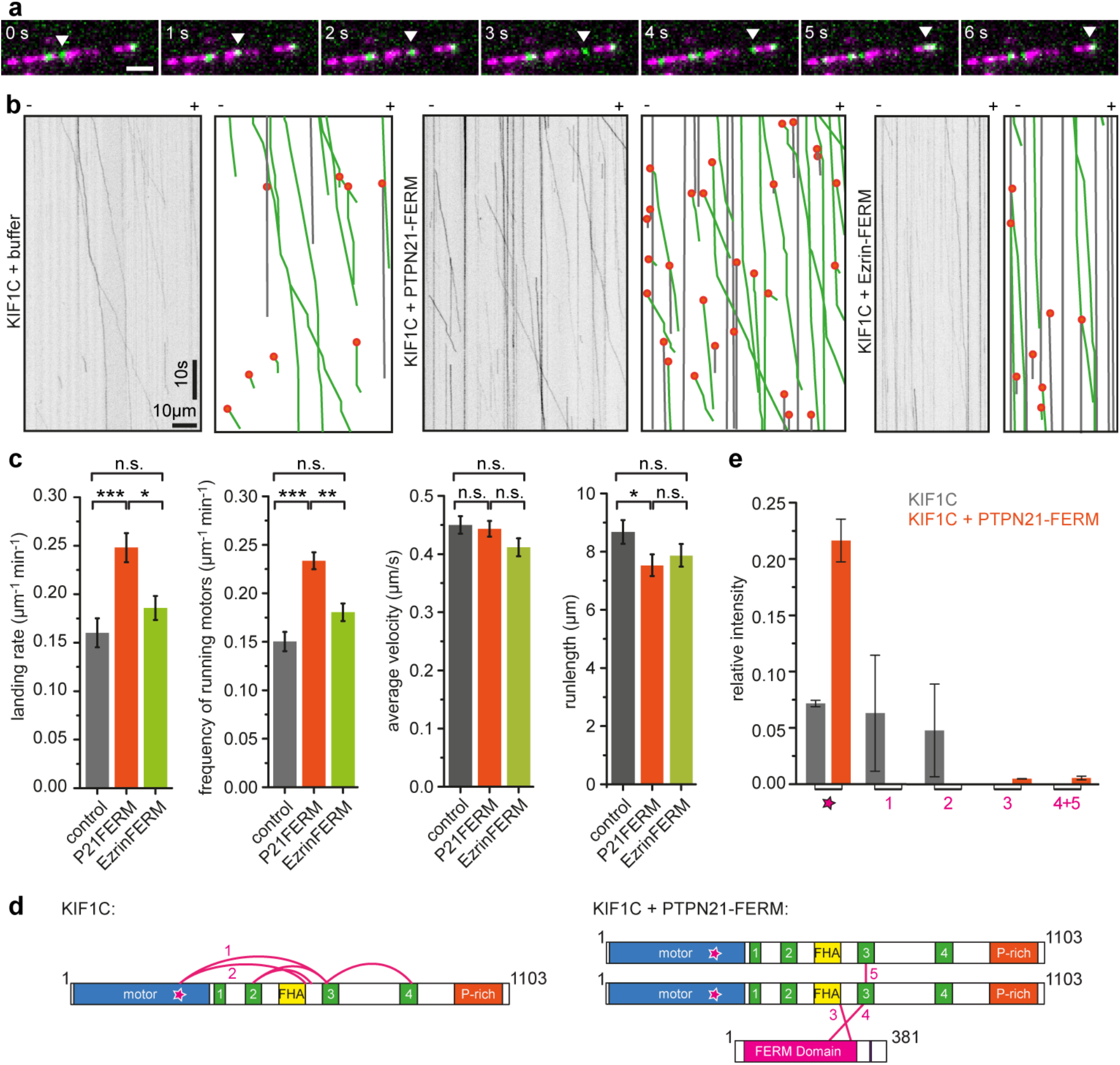
PTPN21 FERM domain activates KIF1C in vitro. **(a)** KIF1C-GFP (green) is a processive motor in single molecule assays on Taxol-stabilised microtubules (magenta). Scale bar 2 μm. **(b)** Representative kymographs from single molecule experiments of KIF1C in the presence of FERM domains of PTPN21 and Ezrin. Grey lines indicate immobile motors, green lines running motors and orange dots landing events. **(c)** Quantification of landing rate, frequency of running motors (>25 nm/s), average velocity and run length. Data show mean±SEM. n= 30, 39 and 21 MTs respectively from three independent experiments. t-test with bonferroni correction: * p<0.05, ** p<0.005, *** p<0.0005, n.s. p>0.05. **(d-e)** Crosslinking mass spectrometry of KIF1C in the presence of PTPN21-FERM identified two crosslinks between KIF1C and the FERM domain and one dimer crosslink in KIF1C (labelled 3,4 and 5). Crosslinks in KIF1C alone are shown for comparison. Star indicates peptide in motor domain that is found in crosslinks with stalk domain (labelled 1 and 2). Relative abundance of un-crosslinked peptide LKEGANINK (star) and the five crosslinked peptides is shown for KIF1C alone and KIF1C + PTPN21-FERM. Data show mean ± SD for three independent experiments. See supplementary figures for ion fragmentation spectra of all crosslinked peptides.

If PTPN21 relieves the autoinhibition, the following two predictions should hold true: (i) PTPN21_FERM_ binding should remove the interactions between stalk and motor domain of KIF1C, and (ii) deleting the region required for auto-inhibition should result in a dominant-active motor that cannot be further activated by binding of PTPN21_FERM_. To test the first prediction, we performed crosslinking mass spectrometry experiments with KIF1C and PTPN21_FERM_. We found a number of new crosslinks, but could no longer find the intramolecular crosslinks in KIF1C (Figure 4d-e). The new crosslinks show interactions of the FERM domain with lysines 591 and 645 in KIF1C as well as an intradimer crosslink between lysines 640 and 645 in KIF1C (Figure S2). Because the crosslinks between motor domain and stalk were absent in the presence of PTPN21_FERM_, the relative abundance of the un-crosslinked LKEGANINK peptide at the microtubule binding interface of KIF1C increased threefold in the KIF1C+ PTPN21_FERM_ sample compared to KIF1C alone (Figure 4e). These data confirm that the presence of the FERM domain is capable of freeing the motor domain, and relieving the autoinhibition of KIF1C. To test the second prediction, we generated a deletion of coiled-coil 3, which makes contact with the motor domain in the auto-inhibited conformation. First, we assessed whether KIF1CΔCC3 is hyperactive by localisation of the construct in cells. We had described previously that KIF1C accumulates specifically in tails of migrating RPE1 cells (Theisen et al., 2012). Expression of KIF1CΔCC3-GFP in RPE1 cells resulted in accumulation of the construct in cell tails and also elsewhere at the periphery of the cell (Figure 5a). As tail accumulation is sensitive to tail state, i.e. whether it is forming or retracting (Theisen et al., 2012), we used KIF1C-mCherry as internal control. In comparison to KIF1C-GFP, which was enriched in tails to a similar extent as KIF1C-mCherry (tail:cytoplasm ratio 6.1±1.3 for KIF1C-mCherry and 7.4±1.2 for KIF1C-GFP, n=26 cells), KIF1CΔCC3-GFP was about 3-fold more enriched (tail:cytoplasm ratio 3.9±0.7 for KIF1C-mCherry and 13.5±2.6 for KIF1CΔCC3-GFP, n=32 cells) (Figure 5b). Secondly, we purified recombinant KIF1CΔCC3-GFP from insect cells and performed single molecule assays. KIF1CΔCC3-GFP had a 3-fold higher landing rate compared to wildtype KIF1C-GFP, and addition of PTPN21_FERM_ did not further increase the landing rate of KIF1CΔCC3-GFP (Figure 5c-d). Therefore, these experiments confirm that binding of PTPN21 to the stalk domain of KIF1C relieves the autoinhibition of KIF1C and thereby enables the motor to engage with microtubules.

**Figure 5:**
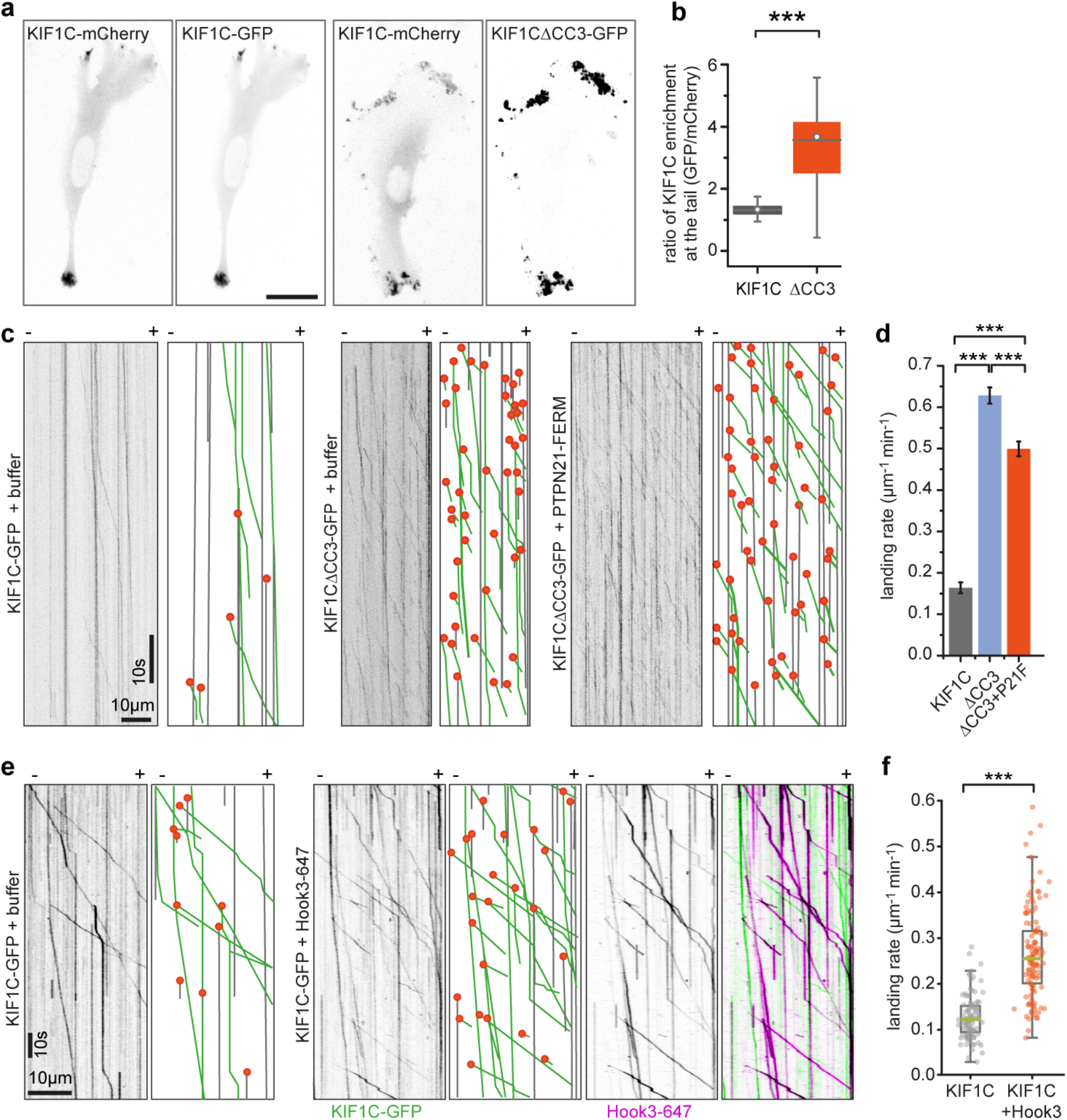
Deletion of coiled-coil 3 in KIF1C stalk results in a hyperactive motor. **(a)** RPE cells cotransfected with full-length KIF1C-mCherry and either KIF1C-GFP or KF1C△CC3-GFP and imaged 36 hours post transfection. Scale bar is 25 μm. **(b)** Box plot shows ratio of KIF1C enrichment at the tail relative to cytoplasmic levels for GFP versus mCherry channel. *** p<0.0005 (Kolmogorov-Smirnov). n=26, 32 cells. **(c)** Representative kymographs from single molecule experiments. Grey lines indicate immobile motors, green lines running motors and orange dots landing events. **(d)** Frequency of KIF1C landing events shown as mean ± SEM from 17, 82 and 58 microtubules, respectively, pooled from three experiments. *** p<0.0005 (t-test with bonferroni correction). **(e)** Representative kymographs from single molecule experiments of KIF1C-GFP alone or in the presence of Hook3 labelled with Alexa647. Motility and landing events indicated as in c. **(f)** Quantification of absolute frequency of KIF1C-GFP motors landing on microtubules, n=116 and 131 MTs, respectively, pooled from three independent experiments. *** p<0.0005 (two-tailed t-test).

### Hook3 binds to same region in KIF1C stalk as PTPN21 and also activates KIF1C

Finally, we wondered how universal this mechanism is and whether other proteins binding KIF1C stalk could activate KIF1C in a similar manner. To identify KIF1C stalk interactors we performed a BioID experiment with KIF1C full length and a deletion construct KIF1CΔ623-825 that lacked both coiled-coil 3 and the previously identified minimal PTPN21 binding site. We identified 240 proteins that were isolated with steptavidin-beads upon biotinylation with KIF1C-BioID2, but were absent from the sample with KIF1CΔ623-825-BioID2. The top hit with most peptides identified was Hook3. Hook3 has previously been identified as an activator of dynein/dynactin - an activity requiring its N-terminal globular domain and all three coiled-coil domains (Olenick et al., 2016, Urnavicius et al., 2018, Schroeder and Vale, 2016, McKenney et al., 2014). In contrast, the C-terminus of Hook3 was shown to interact with KIF1C (Redwine et al., 2017). To test whether Hook3 could also activate KIF1C, we purified recombinant human Hook3 with a C-terminal SNAP-tag from insect cells, labelled it with Alexa647 and performed single molecule assays with KIF1C. In the presence of Hook3, the landing rate of KIF1C increased 2-fold and we observed co-transport of Hook3-Alexa647 with KIF1C-GFP (Figure 5e-f), thus confirming that Hook3 functions as an activator similarly to PTPN21 by binding to the KIF1C stalk region adjacent to the site engaged in intramolecular interactions of KIF1C during autoinhibition.

## DISCUSSION

Our findings support a model whereby KIF1C is a stable dimer that is held in an autoinhibited conformation by interaction of its stalk region including the third coiled-coil domain with the microtubule binding surface of the motor domain (Figure 6a). Autoinhibition is relieved upon binding of PTPN21 FERM domain or Hook3, allowing the motor domain to engage with microtubules (Figure 6a-b). While the model agrees with the finding that KIF1C motors are dimeric in cells (Dorner et al., 1999), this is in contrast to the mode of autoinhibition described for other kinesin-3 motors, KIF1A, Unc104, KIF16B, KIF13A and KIF13B that all undergo a monomer-dimer transition (Al-Bassam et al., 2003, Soppina et al., 2014, Tomishige et al., 2002, Rashid et al., 2005). However, interactions of the stalk or tail region with the motor domain have also been described for KIF13B and KIF16B (Farkhondeh et al., 2015, Yoshimura et al., 2010). These might act as a second layer of activity control once dimers are formed or stabilise the inhibited monomer conformation. Our data suggest that KIF1C autoinhibition acts by steric blockage of the microtubule binding. In contrast, kinesin-1, which is also autoregulated by a tail-block mechanism, is inhibited by crosslinking of the motor domains that prevent movement required for neck linker undocking (Kaan et al., 2011, Stock et al., 1999).

**Figure 6:**
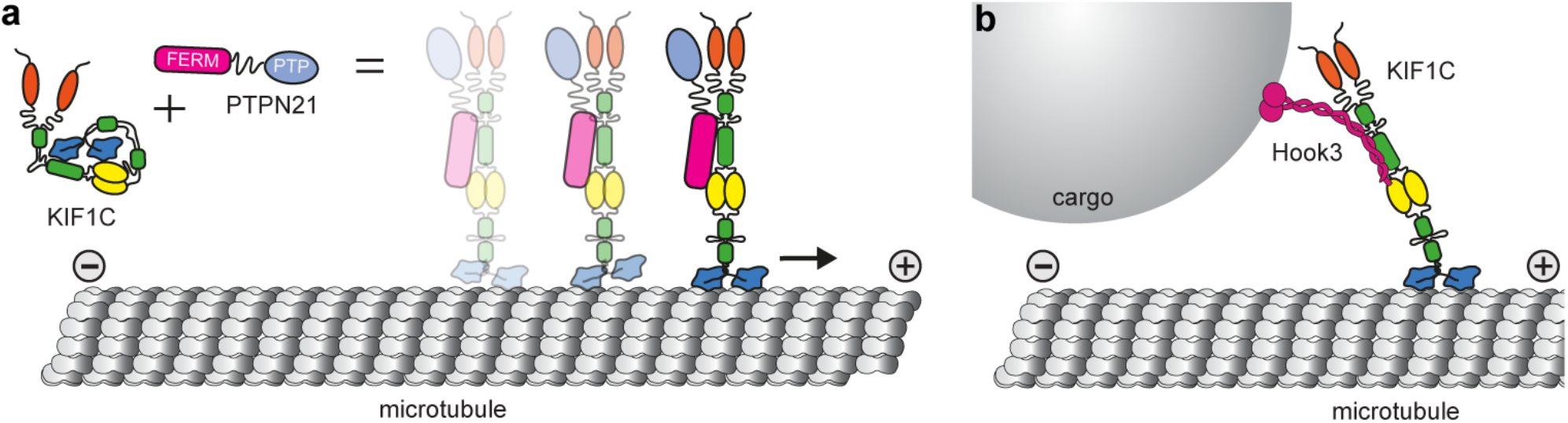
Activation of KIF1C. **(a)** KIF1C is an autoinhibited dimer in solution. Intramolecular interactions between stalk and motor domain prevent the inhibited motor from interacting with microtubules. Upon binding of PTPN21, the intramolecular interactions are released, KIF1C can engage with microtubules and will step processively towards the plus end. **(b)** The cargo adapter Hook3 also binds to the stalk of KIF1C and activates it, thereby mediating cargo-activated transport along microtubules.

We identify the phosphatase PTPN21 as a KIF1C activator. The first 378 amino acids containing the FERM domain are sufficient to activate KIF1C *in vitro* and in cells and does neither require the catalytic activity nor the phosphatase domain of PTPN21. Even though a scaffold activity is sufficient, it is still possible that PTPN21-mediated dephosphorylation of KIF1C (Dorner et al., 1998) can further modulate the activity of KIF1C, modify KIF1C cargo, cargo adapters or the activity of adjacent motors, a possibility that will be interesting to explore in a future study. PTPN21 has been shown to dynamically localise to focal adhesions (Carlucci et al., 2008) or EGF receptor recycling sites (Roda-Navarro and Bastiaens, 2014). It is possible that KIF1C-mediated transport facilitates the efficient turnover of PTPN21 at these sites. Thus the phosphatase could be both a cargo and a regulator of KIF1C. We have implicated KIF1C previously in the transport of integrins required for the maturation of trailing focal adhesions (Theisen et al., 2012). It is possible that additionally, KIF1C transport facilitates PTPN21-mediated regulation of Src tyrosine kinase and FAK signalling that promote cell adhesion and migration (Carlucci et al., 2008).

PTPN21 FERM was able to compensate for partial depletion of KIF1C, probably by activating the remaining KIF1C motors, but it is also possible that PTPN21 activates additional kinesins which can be recruited to the cargoes studied here. The most likely candidate for this is KIF16B, a highly processive early endosome transporter that has been shown to also interact with PTPN21 FERM (Carlucci et al., 2010, Hoepfner et al., 2005, Soppina et al., 2014). The expression of the FERM domain efficiently stimulated dense core vesicle transport in primary neurons. PTPN21 has been associated with schizophrenia in a genome-wide association study (Chen et al., 2011) and shown to promote neuronal survival and growth (Plani-Lam et al., 2015, Cui et al., 2017). While the latter is thought to occur via NRG3 or ERK1/2 signalling, neuronal function and morphology is compromised when microtubule transport is perturbed and the function of PTPN21 as neuronal transport regulator might contribute to the complications in schizophrenia patients with PTPN21 mutations. Further, cAMP/PKA pathway stimulation of kinesin-1 has been shown to reverse aging defects in *Drosophila* neurons, and a similar age-driven decrease in KIF1C transport of dense-core vesicles and other organelles may have similar effects in neurodegenerative diseases (Vagnoni and Bullock, 2018). The findings that Hook3 can activate both dynein/dynactin (McKenney et al., 2014, Olenick et al., 2016, Schroeder and Vale, 2016) and KIF1C (this study) and that the binding sites for these opposite directionality motors are non-overlapping (Redwine et al., 2017, Schroeder and Vale, 2016), suggests that Hook3 could simultaneously bind to KIF1C and dynein/dynactin and provide a scaffold for bidirectional cargo transport. How the directional switching would be orchestrated in such a KIF1C-DDH complex is an exciting question for the future. It is important to note that Hook3 is not the only dynein cargo adapter which binds KIF1C. BICDR1 has been shown to bind to the proline-rich C-terminal region of KIF1C (Schlager et al., 2010), and BICD2 appears to interact with KIF1C biochemically (Novarino et al., 2014). Whether BICDR1 or BICD2 are able to activate the motor is unclear, but it is possible that different adapters not only mediate linkage to a different set of cargoes, but also recruit opposite polarity motors in different conformations and thus relative activity. For dynein/dynactin, such a difference is seen in BICD2 recruiting only one pair of dynein heavy chains while BICDR1 and Hook3 recruit two pairs and thus are able to exert higher forces (Urnavicius et al., 2018). BICDR1 also binds Rab6 and recruits both dynein/dynactin and KIF1C to participate in the transport of secretory vesicles (Schlager et al., 2010). Rab6 in turn has been shown to bind and inhibit the KIF1C motor domain (Lee et al., 2015). This could provide a potential mechanism for a second layer of regulatory control of KIF1C activity to facilitate its minus end-directed transport with dynein-dynactin-Hook3.

Taken together, we provide mechanistic insight into the regulation of KIF1C, a fast long-distance neuronal transporter. We show that KIF1C is activated by a scaffold function of PTPN21 and the dynein cargo adapter Hook3, but the mechanism of autoinhibition release described here is likely to be universal and we expect cargoes and further scaffold proteins binding to the stalk region around the third coiled-coil in KIF1C to also activate the motor and initiate transport along microtubules. This opens up new research avenues into how KIF1C activity is controlled in space and time in cells.

## METHODS

### Plasmids and siRNAs

The following plasmids used in this study were described previously: *pKIF1C^RIP2^GFP* (Efimova et al., 2014), *pKIF1C-mCherry* (Theisen et al., 2012), *pFlag* (Theisen et al., 2012, Efimova et al., 2014), *pα5-integrin-GFP* (Laukaitis et al., 2001), *pNPY-RFP* (Miller et al., 2009), *pHA-PTPN21-WT* and *pHA-PTPN21-C1108S* (Cardone et al., 2004).

*pFastBac-M13-6His-KIF1CGFP* was generated by digesting *pKIF1C^RIP2^GFP* with *Eco*RI - *Mfe*I and replacing it in the backbone of *pFastBac-M13* (Invitrogen). A resulting frameshift between the 6His tag and the N-terminus of KIF1C was corrected by cutting with *Eco*RI, mung bean nuclease treatment and religation of the vector. *pKIF1CΔCC3-GFP* was generated by introducing a *Sal*I restriction site after amino acid position D679 in pKIF1C^RIP2^GFP by PCR using primers 5’- CGGGgtcgacTCTGACAAGCGCTCTTG-3’ and 5’-GCCGTTTACGTCGCCGTC-3’. The *Sal*I-*BamH*I fragment was replaced in *pKIF1C^RIP2^GFP* with the truncated fragment to create a plasmid expressing KIF1C^RIP2^Δ623_679-GFP in mammalian cells. *pKIF1C△623-825^RIP2^GFP* was generated by introducing a *SalI* restriction site after amino acid position D825 in pKIF1C^RIP2^GFP by PCR using primers 5’-GGAGgtcgacCGAGGGGCGGAGGTGG-3’ and 5’-GCCGTTTACGTCGCCGTC-3’ and introduced as SalI-BamHI fragment to *pKIF1C^RIP2^GFP as above*.

*pFastBac-M13-6His-KIF1CΔCC3-GFP* was generated by digesting *pKIF1CΔCC3-GFP* with *Bsi*WI-*Bam*HI and replacing it in the backbone of *pFastBac-M13-6His-KIF1CGFP*.

RNAi-protected PTPN21 was made using *pHA-PTPN21-WT* as a template in a three-step mutagenesis PCR with upstream primer 5’-GAGCCCTCTGTATTTCTGATG-3’, downstream primer 5’-CGCAAATGGGCGGTAGGCGTG-3’ and mutagenesis primer 5’-GTTGTAGTACCAtAatgaaAAGTAAGTGACC-3. The fragment containing the mutation was replaced in *pHA-PTPN21-WT* using *Nhe*I to generate *pHA-PTPN21^RIP^*. RNAi protected inactive PTPN21, *pHA-PTPN21^RIP^C1108S* was generated by replacing the fragment containing mutation C1108S in *pHA-PTPN21^RIP^* using BsgI.

*pFERM* expressing HA-tagged PTPN21_1-377_ was generated by digesting *pHA-PTPN21^RIP^* and *pEGFP-N1* (Clonetech) with *Hind*III and inserting the fragment in *pEGFP-N1*. As the GFP is not in frame, the FERM domain is expressed from this plasmid with an N-terminal HA-tag only.

To express and purify FERM domains from *E. coli, pET22b-HA-PTPN21_1-381_-6His* was generated by amplifying *pHA-PTPN21^RIP^* by PCR with primers 5’-AAGCTTcatATGGGATACCCATACGATGTTC-3’ and 5’-GAGGTCAgcggccgcTCTATCCAAGCTTGTCTG-3’, digesting with *Nde*I and *Not*I and replacing in *pET22b-mEB1-6His* (Roth et al., 2018). FERM domain from Ezrin, *pET22b-HA-Ezrin_1-328_-6His* was generated by PCR from random-primed cDNA reverse transcribed from RPE1 RNA using primers 5’-GAAACCcatatgGGATACCCATACGATGTTCCAGATTACGCTGTGGTGCCGAAACCAATCAATGT C-3’ and 5’-CGGTTgcggccgcTTTCTTCTCTGTTTCCAGCTG-3’, digesting with *Nde*I and *Not*I and replacing in *pET22b-mEB1-6His*.

A multi-tag version of *pFastBacM13* (Invitrogen) was created by ligating an insect-cell codon optimised synthesised cDNA for 8His-ZZ-LTLT-BICDR1-SNAPf between the *RsrI*I and *Not*I sites creating *pFastBacM13-8His-ZZ-LTLT-BICDR1-SNAPf*. Hook3 was cloned into this by PCR from random-primed cDNA reverse transcribed from RPE1 RNA using primers which incorporated a 5’ AscI and a 3’ *BamH*I site allowing direct ligation into the multi-tag *pFastBacM13* vector creating *pFastBacM13-8His-ZZ-LTLT-Hook3-SNAPf*, which was subsequently used for production of a recombinant baculovirus. The forward and reverse primers were: 5’-ACATAggcgcgccCTATGTTCAGCGTAGAGTC-GCTG-3’ and 5’-ACAATAACCGGTggatccCTTGCTGTGGCCGGCTG-3’.

Human codon-optimised BioID2-HA (Kim et al., 2016) was synthesised to include 5’ BamHI and 3’ NotI restriction sites allowing direct substitution of the C-terminal GFP of *pKIF1C^RIP2^GFP* and *pKIF1C△623-825^RIP2^GFP*with a BioID2-HA tag to generate *pKIF1C^RIP2^-BioID2-HA* and *pKIF1C△623-825^RIP2^-BioID2-HA*.

siRNA oligonucleotides for KIF1C and PTPN21 were: siControl 5’-GGACCUGGAGGUCUGCUGU-[dT]-[dT]-3’ (Theisen et al., 2012), siKIF1C-2 (targeting both rat and human KIF1C) 5’-GUGAGCUAUAUGGAAAUGU-[dA]-[dC]-3’ (Theisen et al., 2012, Efimova et al., 2014), siPTPN21 (targeting both rat and human PTPN21) 5’-UUCAGCCUCUGGUACUACA-[dT]-[dT]-3’.

### Recombinant protein purification

For baculovirus expression, *pFastBac-M13-6His-KIF1CGFP, pFastBac-M13-6His-KIF1CΔCC3-GFP, pFastBacM13-8His-ZZ-LTLT-Hook3-SNAPf* plasmids were transformed into DH10BacYFP competent cells and plated on LB-Agar supplemented with 30 μg/ml kanamycin (#K4000, Sigma), 7 μg/ml gentamycin (#G1372, Sigma), 10 μg/ml tetracycline (#T3258, Sigma, 40 μg/ml Isopropyl β-D-1-thiogalactopyranoside (IPTG, #MB1008, Melford) and 100 μg/ml X-Gal (#MB1001, Melford). Positive transformants (white colonies) were screened by PCR using M13 forward and reverse primers for the integration into the viral genome. The bacmid DNA was isolated from the positive transformants by the alkaline lysis method and transfected into SF9 cells (Invitrogen) with Escort IV (#L-3287, Sigma) according to manufacturer’s protocols. After 5-7 days, the virus (passage 1, P1) is harvested by centrifugation at 300×g for 5 min in a swing out 5804 S-4-72 rotor (Eppendorf). Baculovirus infected insect cell (BIIC) stocks were made by infecting SF9 cells with P1 virus and freezing cells before lysis (typically around 36 hr) in a 1° cooling rate rack (#NU200 Nalgene) at -80°C. P1 virus was propagated to passage 2 (P2) by infecting 50 ml of SF9 culture and harvesting after 5-7 days as described above (Wasilko et al., 2009). For large scale expression, 500 ml of SF9 cells at a density of 1-1.5×10^6^ cells/ml were infected with one vial of BIIC or P2 virus. Cells were harvested when 90% infection rate was achieved as observed by YFP fluorescence, typically between 48-72 hr. Cells were pelleted at 252 ×g in a SLA-3000 rotor (Thermo Scientific) for 20 min. The pellet was resuspended in 4 ml of SF9 lysis buffer (50 mM Sodium phosphate pH 7.5, 150 mM NaCl, 20 mM Imidazole, 0.1% Tween 20, 1.5 mM MgCl_2_) per gram of cell pellet, supplemented with 0.1 mM ATP and cOmplete protease inhibitor cocktail (#05056489001, Roche).

Bacterial expression plasmids, *pET22b-HA-PTPN21_1-381_-6His* and *pET22b-HA-Ezrin_1-328_-6His* were transformed into BL21 DE3 (Invitrogen) and single colonies were grown overnight as 3 ml starter cultures. The starter cultures were diluted 1:100 and grown in 200 ml of 2×YT (yeast extract, tryptone 16g) at 37°C, 180 rpm until they reached an OD_600nm_ of 0.5. Expression was induced with 500 μM IPTG and incubated at 16°C overnight. Cells were harvested at 1500×g in a SLA-3000 rotor and resuspended in bacterial lysis buffer (50 mM Sodium phosphate pH 7.5, 50 mM NaCl, 20 mM Imidazole, 1.5 mM MgCl_2_) supplemented with 1 mM phenylmethanesulfonyl fluoride (PMSF) (#MB2001; Melford).

Purification of recombinant KIF1C-GFP, KIF1CΔCC3-GFP, PTPN21-FERM and Ezrin-FERM was carried out in a two-step process at 4°C. Harvested cell pellets were lysed using a douncer (#885301, Kontes) with 20 strokes. Bacterial cell pellets were sonicated at 50% amplitude for 30s in a 10s on off cycle repeated thrice. Lysates were then cleared by centrifugation at 38,000 ×g in a SS-34 rotor (Sorvall) for 30min. SP Sepharose beads (#17-0729-01, GE Healthcare) were equilibrated with the lysis buffer and the cleared lysate obtained is mixed with the beads and batch bound for 1 hour. Next, the beads were loaded onto a 5ml disposable polypropylene gravity column (#29922, Thermo Scientific) and washed with 10 CV SP wash buffer (50 mM Sodium phosphate pH 7.5, 150 mM NaCl) and eluted with SP elution buffer (50 mM Sodium phosphate pH 7.5, 300 mM NaCl). The peak fractions obtained were pooled and diluted with Ni-NTA lysis buffer (50 mM Sodium phosphate pH 7.5, 150 mM NaCl, 20 mM Imidazole, 10% glycerol) and batch bound to Ni-NTA beads (#30230, Qiagen) for 1 hour. The beads were loaded onto a gravity column and washed with 10 CV of Ni-NTA wash buffer (50 mM Sodium phosphate pH 7.5, 150 mM NaCl, 50 mM Imidazole and 10% glycerol) and eluted with Ni-NTA elution buffer (50 mM Sodium phosphate pH 7.5, 150 mM NaCl, 150 mM Imidazole, 0.1 mM ATP and 10% glycerol). The peak fractions were run on a SDS-PAGE gel for visualisation and protein was aliquoted, flash frozen and stored in liquid nitrogen.

Hook3-647 was purified and labelled in a two-step process utilising both the His and ZZ affinity tags at 4°C. A pellet corresponding to 500 ml insect culture was resuspended in 40 ml Adaptor Lysis Buffer (50 mM HEPES pH 7.2, 150 mM NaCl, 20 mM imidazole, 1×cOmplete protease inhibitor cocktail, 1mM DTT) and lysed with 20 strokes in a douncer. Lysates were cleared of insoluble material by centrifugation at 50,000 ×g for 40 minutes in a Sorvall T-865 rotor. The cleared lysate was bound to 2 ml Ni-NTA beads for 2 hours, and subsequently washed with 200 CV of lysis buffer followed by 200 CV lysis buffer supplemented with 60 mM imidazole. Protein was eluted in 5 CV of lysis buffer containing 350mM Imidazole, and the eluate was batch-bound to 1ml IgG Sepharose (#17-0969-01, GE Healthcare Life Sciences) for 2 hours. Beads were briefly washed with TEV cleavage buffer (50 mM Tris–HCl pH 7.4, 148 mM KAc, 2 mM MgAc, 1 mM EGTA, 10% (v/v) glycerol, 1 mM DTT) before being collected in a 1.5ml Eppendorf tube and incubated with 3.5 μM SNAP-Surface Alexa Fluor 647 substrate (#S9136S, New England Biolabs) for 2 hours. Beads were washed with 200 CV TEV cleavage buffer and resuspended in a 1.5 ml Eppendorf tube containing 50 μg/ml TEV protease, which was allowed to digest the protein off of beads overnight at 4°C. Finally, eluate was collected from the beads, concentrated to between 5-10μM in a Centrisart I 20KDa ultrafiltration device (#13249-E, Sartorius), flash frozen and stored in liquid nitrogen.

### Hydrodynamic analysis

Size exclusion chromatography was carried out using Superdex 16/60 200pg (#28989335, GE Healthcare) column on an AKTApurifier10 FPLC controlled by UNICORN 5.11 (GE Healthcare). The column was equilibrated with the SEC Buffer (35 mM Sodium Phosphate pH 7.5, 150 mM NaCl, 1.5mM MgCl_2_). 200μl of ~0.5 mg/ml KIF1C-GFP was injected into the column and 0.5 ml fractions were collected using the fraction collector. GFP fluorescence in the fractions was measured using Nanodrop 3300 Fluorospectrometer (Thermo Scientific), to determine peak positions. 100 μl of 5 mg/ml standard proteins were injected individually. The Stokes radius for the standard proteins used were as follows thyroglobulin (#T9145, Sigma) 8.5 nm (Edelhoch, 1960), apoferritin (#A3660, Sigma) 6.1 nm (Björk, 1973), catalase (#C9322, Sigma) 5.2 nm (Sund et al., 1967), bovine serum albumin (#A7906, Sigma) 3.48 nm (Creeth, 1952). Log R_s_ versus (elution volume-void volume) for standard proteins was plotted and stokes radius R_s_ for KIF1C was determined from the linear fit: y = (0.973±0.04) – (0.014±0.002)•x. Three independent repeats were carried out at 500 mM NaCl and two at 150 mM NaCl.

5ml of 10 - 40% vol/vol glycerol gradients were made using the Gradient master (#108, Biocomp) in 35 mM Sodium phosphate pH 7.5, 1.5 mM MgCl_2_, 0.1 mM ATP, 1 mM EGTA at different NaCl concentration (50, 150, 250, 350, 500 mM NaCl). 100 μl of ~0.25 mg/ml KIF1C-GFP was loaded on top of the gradient and the samples were centrifuged at 364,496 ×g in SW55Ti (Beckman Coulter) for 16 hrs at 4°C. Gradients were fractionated by carefully pipetting 200 μl aliquots from the top. Absorbance at 280nm was measured in Nanodrop 2000c Spectrophotometer and fluorescence using Nanodrop 3300 Fluorospectrometer (Thermo Scientific) to determine peak positions. 100 μl of 5mg/ml standard proteins were loaded individually on separate gradients. The samples were processed for SDS-PAGE and immunoblotting with anti-KIF1C primary (1:4000, #AKIN11, Cytoskeleton) and anti-rabbit IgG HRP conjugate secondary (1:4000, #W401B Promega) antibodies. The sedimentation co-efficient of standard proteins were: apoferritin 17.6 S (Rothen, 1944), catalase 11.3 S (Sund et al., 1967), glucose oxidase (#G7141, Sigma) 8 S (Tsuge et al., 1975), bovine serum albumin 4.6 S (Creeth, 1952). Sedimentation co-efficient versus elution volume for standard proteins was plotted and sedimentation co-efficient for KIF1C at different salt concentration was calculated from the linear fit: y = (5.57±0.23)•x – (3.84±0.63). Independent repeats were carried out thrice (500 mM NaCl and 50 mM NaCl), twice (150 mM and 250 mM) and once (350 mM) respectively.

The molecular weight was calculated from stokes radius and sedimentation co-efficient using equation M_r_ = 4205•Rs•S as described in(Erickson, 2009). The frictional co-efficient was calculated from the following formula, *f/f_min_* = *S_max_/S* with *S_max_* = 0.00361 •*M*^2/3^ using M=308,000 Da for KIF1C-GFP dimer and S is the sedimentation co-efficient determined from glycerol gradient centrifugation (Erickson, 2009).

### Crosslinking mass-spectrometry

Two cross-linkers, BS3 (bis (sulpho-succinimidyl) suberate) (#21580, Thermo Scientific) and EDC (1-Ethyl-3-(3-dimethylaminopropyl) carbodiimide) (#22980, Thermo Scientific) were used to analyse protein interactions using crosslink mass-spectrometry. 5 mM of cross-linker was freshly prepared in MilliQ water and was mixed in the ratio of 1:2 by pipetting with the protein of interest (1 μM) in solution. KIF1C-GFP and PTPN21-FERM were mixed in equimolar amounts. For EDC cross-linking, 3 mM N-hydroxysulfosuccinimide (#24510, Thermo Scientific) was added to the reaction with EDC to improve efficiency of the cross-linking. The reaction was incubated at room temperature for an hour shaking at 400 rpm and quenched with 50 mM Tris HCl pH 7.5. Next, the protein was diluted in equal volume of 50 mM Ammonium bicarbonate (#A6141, Sigma) and reduced using 1 mM DTT (#MB1015, Melford) for 60 min at room temperature. The sample was then alkylated with 5.5 mM iodoacetamide (#I6709, Sigma) for 20 min in the dark at room temperature and digested using 1 μg trypsin (sequencing grade; #V5111, Promega) per 100 μg of protein overnight at 37°C. The crosslinked peptides were de-salted using C18 stage tips. 20 μl samples were then analysed by nano LC-ESI-MS/MS using UltiMate^®^ 3000 HPLC series for peptide concentration and separation. Samples were separated in Nano Series™ Standard Columns. A linear gradient from 4% to 35% solvent B (0.1% formic acid in acetonitrile) was applied over 30 min, followed by a step change 35% to 70% solvent B for 20 min and 70% to 90% solvent B for 30 min. Peptides were directly eluted (~ 300 nL min^-1^) via a Triversa Nanomate nanospray source into a Orbitrap Fusion mass spectrometer (Thermo Scientific). Positive ion survey scans of peptide precursors from 375 to 1500 *m/z* were performed at 120 K resolution (at 200 *m/z)* with automatic gain control 4x10^5^. Precursor ions with charge state 2–7 were isolated and subjected to HCD fragmentation in the Orbitrap at 30 K resolution or the Ion trap at 120 K. MS/MS analysis was performed using collision energy of 33%, automatic gain control 1x10^4^ and max injection time of 200 ms. The dynamic exclusion duration was set to 40 s with a 10 ppm tolerance for the selected precursor and its isotopes. Monoisotopic precursor selection was turned on. The instrument was run in top speed mode with 2 s cycles.

Raw data files were converted to .mgf format using the ProteoWizard msconvert toolkit (Chambers et al., 2012). Sequences are visualized using Scaffold (Proteome software) for percentage coverage and purity followed by analysis using StavroX(Gotze et al., 2015). Cross-linked peptides were identified using the StavroX software with appropriately defined parameters for the cross-linker used. A cross-linked peptide was considered as valid if it achieved a StavroX score of at least 100, which is based on: (i) the presence of ion series fragmentation for both peptides, specifically those fragment ions that include the cross-linker and the attached second peptide, (ii) the proximity of observed fragmentation ion mass to expected fragment ion mass, (iii) number of ion fragments for b or y ions in the a peptide, (iv) number of unidentified high intensity signals in the spectra. The MS/MS spectra were also manually inspected and only those crosslinks were accepted for which fragmentation ions were observed for both peptides and 3 or more fragments for b or y ions in the alpha peptide were required to match. Any crosslinks of continuous peptides, which could indicate intradimer interactions were rejected as these could not be distinguished from partially cleaved peptides that have been modified by the crosslinker. The relative abundance of peptides was determined using MaxQuant label-free quantitation of crosslinked precursor ions. The integrated peak areas were normalised to the three most abundant uncrosslinked peptides for each samples. BS3 experiments were repeated three times and EDC experiments twice for each sample.

### BioID protein interaction analysis

For each BioID experiment, three 14.5cm dishes of RPE1 cells were grown, and each was transfected at 90% confluency with 20μg DNA diluted in 1ml Optimem (#<u>31985062</u>, Fisher) with 60μg polyethylenimine (PEI, #408727, Sigma). After 24 hours, media was replaced and supplemented with 50μM biotin (#8.51209, Sigma). After a further 12 hours, cells were harvested by trypsinisation, washed with PBS and lysed in RIPA buffer (#9806S, New England Biolabs) supplemented with 250U of benzonase (#E1014, Sigma). Lysate was cleared by centrifugation in a tabletop centrifuge at 20,238xg for 30 minutes at 4°C (#5417R, Eppendorf), and bound to 50μl streptavidin-coated Dynabeads (#65601, Invitrogen) for 1.5 hours, washing away unbound proteins with 3×1ml of PBS. Beads were resuspended in 45μl 50mM ammonium bicarbonate, reduced by adjusting to a final concentration of 10mM TCEP, and subsequently alkylated by adjusting to 40mM chloroacetamide (#C0267, Sigma), incubating at 70°C for 5 minutes. Proteins were digested off of the beads with 0.5μg Trypsin (#V5111, Promega) at 37°C overnight. Peptides were separated from beads by filtration through a 0.22μm tube filter (#CLS8169, Sigma) and desalted using C18 stage tips. 20 μl of peptides were then analysed by nano LC-ESI-MS/MS in an Ultimate 3000/Orbitrap Fusion. Hits were automatically identified, thresholded and viewed in Scaffolds software.

### Cell culture, siRNA and DNA transfections, and drug treatments

A7r5 rat vascular smooth muscle cells were grown and maintained as described previously (Efimova et al., 2014). For siRNA oligonucleotide transfection, HiPerFect (#301704 Qiagen) was used according to manufacturer’s protocol. Rescue DNA plasmids were transfected using Fugene6 (#E2691, Promega) 24 hr after siRNA transfection, cells seeded onto 16 mm glass coverslips (#1232-3148 Fisherbrand) coated for 24 hr with 10 μg/ml fibronectin (#F1141 Sigma) and analysed 72 hr after siRNA transfection. Podosome formation was stimulated in A7r5 cells by treatment with 5 μM PDBu (Phorbol 12,13-dibutyrate; #P1269 Sigma) for 1 hour. For inhibition of Myosin IIA contractile activity, Y27632 Rock inhibitor (#Y0503 Sigma) or Blebbistatin (#B0560 Sigma) were added at various concentrations as indicated in figure labels at the same time as PDBu. DMSO (#D2438, Sigma) was used as negative control.

The RPE1 α5-integrin-GFP stable cell line was established as follows: hTERT RPE1 cells (Clontech) were transfected with pα5-integrin-GFP (Laukaitis et al., 2001) followed by selection with 500 μg/ml Geneticin (#G8168 Sigma) and expansion of single colonies that were screened using fluorescence microscopy. The α5-integrin-GFP cell line was grown in RPE1 growth medium (DMEM/Nutrient F-12 Ham (#D6421 Sigma), 10% FBS (Sigma), 2 mM L-Glutamine (Sigma), 100 U/ml Penicillin (Sigma), 100 μg/ml Streptomycin (Sigma) and 2.3 g/l Sodium Bicarbonate (#S8761 Sigma)) supplemented with 500 μg/ml Geneticin (Sigma). RPE1 cells were transfected with siRNA using Oligofectamine (#12252011 Invitrogen), with DNA using Fugene 6 and analysed 72 hr after siRNA transfection.

Hippocampi were dissected from P1 or P2 mice brains and incubated with papain (#P4762 Sigma) in cold dissection medium (Earl’s buffered salt solution EBSS (#14155 Gibco), 10 mM HEPES (#H0887 Sigma) and 100 U/ml pen-strep) for 15 min at 37°C. After digestion, the hippocampi were spun at 300 ×g for 2 min in a swing out 5804 S-4-72 rotor, washed (10 ml) and resuspended (1 ml) in plating media (modified eagle medium (MEM) (#51200-046 Gibco) supplemented with 100 U/ml pen-strep, 1% N2 (#17502-048 Gibco), 10% horse serum (#26050-039 Gibco), 20 mM glucose, 1 mM sodium pyruvate (#S8636 Sigma) and 25 mM HEPES). Cells were plated on laminin (#L2020 Sigma)/ poly-D-lysine (#P0899 Sigma) coated coverslips and fed by replacing 50% of the medium with plating medium according to (Granseth et al., 2006). Neurons were transfected at DIV4 or DIV5 with lipofectamine 2000 (#11668027 Thermo Scientific) according to manufacturer’s protocol and imaged the next day.

### Fixed cell Imaging

Cells were fixed for 15 min with 4% Paraformaldehyde (#15714 Electron Microscopy Sciences) diluted in Cytoskeleton Buffer (10 mM MES pH 6.1, 138 mM KCl, 3 mM MgCl_2_, 2 mM EGTA, 0.32 M Sucrose). Fixed cells were incubated for 2 min with 0.1% Triton X100 (#T8787 Fisher Scientific) diluted in PBS. Coverslips were then washed with PBS and incubated for 30 min with 0.5% BSA diluted in PBS/0.1%Tween (PBST). Cells were stained for an hour at room temperature or overnight at 4°C with 1:100 anti-cortactin 4F11, (#05-180 Millipore) primary antibody diluted in 0.5% BSA-PBST solution. Coverslips were washed with PBST and incubated for 45 min at room temperature with 1:300 anti-mouse IgG 647 conjugate (#A31571, Molecular probes) secondary antibody and 1:1000 Acti-stain 555 phalloidin (#PHDH1-A, Cytoskeleton) diluted in 0.5% BSA in PBST. Nuclei were stained with 5μg/ml DAPI (#D9542, Sigma) for 1 min and coverslips washed with PBST prior to mounting on glass slides with Vectashield (#H1000 Vector Laboratories).

Cells were imaged using a Deltavision Elite Wide-field microscope and Z-stacks of individual cells were acquired using a 40x NA 1.49 objective and a Z-spacing of 0.2 μm.

To determine the number of podosomes formed in each cell, images of the cortactin channel were first transformed in a Z-projection. Z-projection images were then segmented using the ImageProAnalyzer 7 software by applying a threshold and a minimal size filter of 16 pixels (2.58μm^2^). Individual objects identified in the cortactin Z-projection were then visually compared to the actin channel to confirm they are podosomes, removed if the cortactin staining didn’t coincide with an actin spot and podosome clusters were split into individual podosomes. All experiments were repeated 3 times with 30 cells being analysed for each condition in each experiment.

### Live cell imaging

Live cells were imaged using a 60× oil NA 1.4 objective on an Olympus Deltavision microscope (Applied Precision, LLC) equipped with eGFP, mCherry filter sets and a CoolSNAP HQ2 camera (Roper Scientific) under the control of SoftWorx (Applied Precision). The environment was maintained at 37° C and 5% CO_2_ using a stage-top incubator (Tokai Hit) and a weather station (Precision control).

RPE1 α5-integrin-GFP cells were seeded 24 hours before imaging into quadrant glass-bottom dishes (#627975, Greiner) coated with 10μg/ml Fibronectin. Cells with tails were selected and the mid region of the tail was bleached using the 488nm laser. Vesicle movement was imaged for 200 time points at 0.7-1.0s per frame as described previously (Theisen et al., 2012). α5-integrin-GFP trafficking was analysed using ImageJ. Kymographs were generated from parallel 21 pixel wide lines covering the area of the tail. Vesicle movement was extracted manually from all kymographs using the segmented line tool. Vesicles moving less than a total of 1.5μm over the period of imaging were classified as stationary.

Primary mouse hippocampal neurons transfected with pNPY-RFP and either pFlag (as control) or pFERM and imaged at DIV5 or DIV6. Images were acquired with 500ms exposure every 1.5 s for 160 s. The frequency of NPY-positive vesicles passing through a location in the neurite was determined manually from kymographs.

RPE1 cells co-transfected with *pKIF1C-mCherry* and either *pKIF1C-GFP* or *pKIF1CΔCC3-GFP* were imaged 36 hours post transfection. Images were acquired with 500ms exposure in the eGFP channel and 1s exposure in the mCherry channel. To determine the ratio of enrichment at the tail, a region of interest was drawn manually surrounding the accumulation observed and the mean intensity was measured at the tail, in the cytoplasm near the nucleus and the image background outside the cell in both channels. The ratio of KIF1C enrichment at the tail was calculated as I_tail_-I_background_ / I_cytoplasm_-I_background_ for GFP versus mCherry channel.

### Single molecule motility assay

Microtubules were assembled from 8 μl of 3.4 mg/ml unlabelled pig tubulin, 0.2 μl of 1 mg/ml HiLyte Fluor 647 tubulin (#TL670M, Cytoskeleton) and 0.5 μl of 0.5 mg/ml biotin tubulin (#T333P, Cytoskeleton) in MRB80 (80 mM PIPES pH 6.8, 4 mM MgCl_2_, 1 mM EGTA, 1 mM DTT). The mixture was incubated on ice for 5 min before adding 8.5 μl of polymerisation buffer (2×BRB80 buffer plus 20 % (v/v) DMSO and 2 mM MgGTP). Microtubules were polymerized at 37 °C for 30-60 min. The sample was diluted with 100 μl MT-buffer (MRB80 plus 30 μM paclitaxel). Unincorporated tubulin was removed by pelleting microtubules at 20,238 × g for 8.5 min at room temperature, washing the pellet with 100 μl MT-buffer and re-pelleting as before. The microtubule pellet was re-suspended in 50 μl of MT-buffer and stored at RT covered from light for at least half a day (maximum 3 days) before use.

Coverslips (22×22) were cleaned by incubating in 2.3 M hydrochloric acid overnight at 60°C. The next day, coverslips were washed with Millipore water and sonicated at 60°C for 5 min. The wash cycle was repeated 5 times. The coverslips were dried using a Spin Clean (Technical video) and plasma cleaned using Henniker plasma clean (HPT-200) for 3 min. Flow chambers were made using clean glass slides (Menzel Gläser Superfrost Plus, Thermo Scientific) and double-sided sticky tape (Scotch 3M) by placing the cleaned coverslip on the sticky tape creating a 100 μm deep flow chamber. The surface was coated with (0.2 mg/ml) PLL(20)-g[3.5]-PEG(2)/PEG(3.4)-Biotin (50%) (#PLL(20)-g[3.5]-PEG(2)/PEGbi, Susos AG). Biotin-647-microtubules were attached to this surface with streptavidin (0.625 mg/ml) (#S4762 Sigma) and the surface was blocked with κ-casein (1 mg/ml) (#C0406 Sigma).

KIF1C-GFP, PTPN21-FERM and Ezrin-FERM were pre-spun at 100,000 ×g for 10 min in an Airfuge (Beckman Coulter). 6 nM KIF1C-GFP was incubated with either Ni-NTA elution buffer or 1.25 μM PTPN21-FERM or 1.25 μM Ezrin-FERM for 15 min at room temperature. The complex was then diluted 1:10 in the motility mix (MRB80 supplemented with 1 mM ATP, 5 mM phosphocreatine (#P7936, Sigma), 7 U/ml creatine phosphokinase (#C3755 Sigma), 0.2 mg/ml catalase, 0.4 mg/ml glucose oxidase, 4 mM DTT, 50 mM glucose (#G8270, Sigma), 25 mM KCl, 100 μM taxol, 0.2 mg/ml κ-casein) and flown into the chamber. Experiments with KIF1CΔCC3-GFP were performed in the same way. To verify that comparable amount of motors were added to the assay, the fluorescence intensity for both KIF1C-GFP and KIF1CΔCC3-GFP solutions were measured prior to incubation and flowing into the chamber. For KIF1C and Hook3 experiments we replaced MRB80 in the motility mix with TIRF assay buffer (30 mM HEPES-KOH pH 7.2, 5 mM MgSO_4_, 1 mM EGTA, 1 mM DTT) as this has previously been described for Hook3-based dynein motility (Urnavicius et al., 2018) and preincubated 320nM KIF1C with 500nM Hook3-647.

Chambers were observed on an Olympus TIRF system using a 100× NA 1.49 objective, 488 nm and 640 nm laser lines, an ImageEM emCCD camera (Hamamatsu Photonics) under the control of xCellence software (Olympus), an environmental chamber maintained at 25° C (Okolab, Ottaviano, Italy).

For single molecule motility assays with KIF1C-GFP, KIF1CΔCC3-GFP and FERM domains images were acquired at 30% laser power every 100 ms for 180 s at an exposure of 60 ms for 488 nm laser line, whereas for experiments with KIF1C-GFP and Hook3, images were acquired at 280 ms for 250 s at an exposure of 150 ms for 488 nm (30% laser power) and 60 ms for 647 nm (10% laser power) laser lines.

Images were analysed by generating kymographs from each microtubule using the ImageJ plugin by Arne Seitz (https://biop.epfl.ch/TOOL_KYMOGRAPH.html) and manually identifying runs by scoring the kymographs for moving and static motors. The speed and run length was calculated from the length and slope of each run using a custom ImageJ macro. Blind analysis was carried out to determine landing rates and frequency of running motors in all data sets.

### Bleach step analysis

A flow chamber was prepared as described above with biotin-647-microtubules immobilized on the coverslip. 600 pM KIF1C-GFP was flown into a chamber with MRB80, 1 mM AMP-PNP, 100 μM taxol and 0.2 mg/ml κ-casein. Images were acquired every 500 ms for 600 s at an exposure of 400 ms using the 488 nm laser line at 50% laser power. Images were analysed using the Plot Profile function in ImageJ for each individual spot and manually counting bleach steps. A mixed binomial distribution for dimers and tetramers of KIF1C was fitted to the data using the equation below, where P(k) is the probability to find k bleach steps, p is the fraction of active GFP and x is the fraction of tetramers.

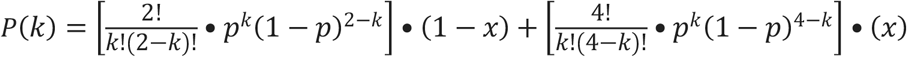

### Statistical Analysis and figure preparation

Statistical data analyses and graphs were generated using Origin Pro 8.5 (OriginLab), python’s Matplotlib and Scipy packages or R. Box plots show quartiles with whiskers indicating 10% and 90% of data. All statistical significance analyses were carried out using two-sample t-tests assuming equal variance, Mann-Whitney U-test or Kolmogorov-Smirnov test. Where necessary, p-values were adjusted for multiple comparisons using Bonferroni and Holm-Šídák corrections. Figures were prepared by adjusting min/max and inverting look up tables using ImageJ and assembled using Adobe Illustrator.

## Supporting information

## ACKNOWLEDGEMENTS

We thank Antonio Feliciello for PTPN21 plasmids, Alex Jones and the WPH Proteomics RTP at the University of Warwick for mass spectrometry, Robert Dallmann for providing mouse brains, Steve Royle for help with hippocampus dissection, and Masanori Mishima for the mixed binomial model and comments on the manuscript.

This work was funded by a non-clinical PhD studentship from the British Heart Foundation (FS/13/42/30377) that supported AB, a Chancellor’s International PhD Scholarship from the University of Warwick that supported NS, and a PhD studentship from the EPSRC-funded MOAC CTD (EP/F500378/1) that supported HH. A.J.Z. is funded by the MRC Doctoral Training Partnership (MR/N014294/1). AS is a Prize Fellow of the Lister Institute of Preventive Medicine and a Wellcome Trust Investigator (200870/Z/16/Z). DR is supported in part from the University of Warwick.

## AUTHOR CONTRIBUTIONS

AS and IK perceived the project, NS purified KIF1C, KIF1CΔCC3 and FERM domains, performed biochemical characterization, single molecule experiments with FERM domain proteins and KIF1CΔCC3, crosslink mass spectrometry, KIF1CΔCC3 localisation in cells and data analysis. AB performed all cell biology experiments, AJZ performed BioID, purified Hook3 and performed single molecule experiments and data analysis, HH analysed crosslink mass spectrometry data, DR generated resources and performed image analysis, AS managed the project and wrote the manuscript with contributions from all authors.

The authors declare no competing interests.

