## Supplementary material for "Activation of intracellular transport by relieving KIF1C autoinhibition"

### SUPPLEMENTARY FIGURES

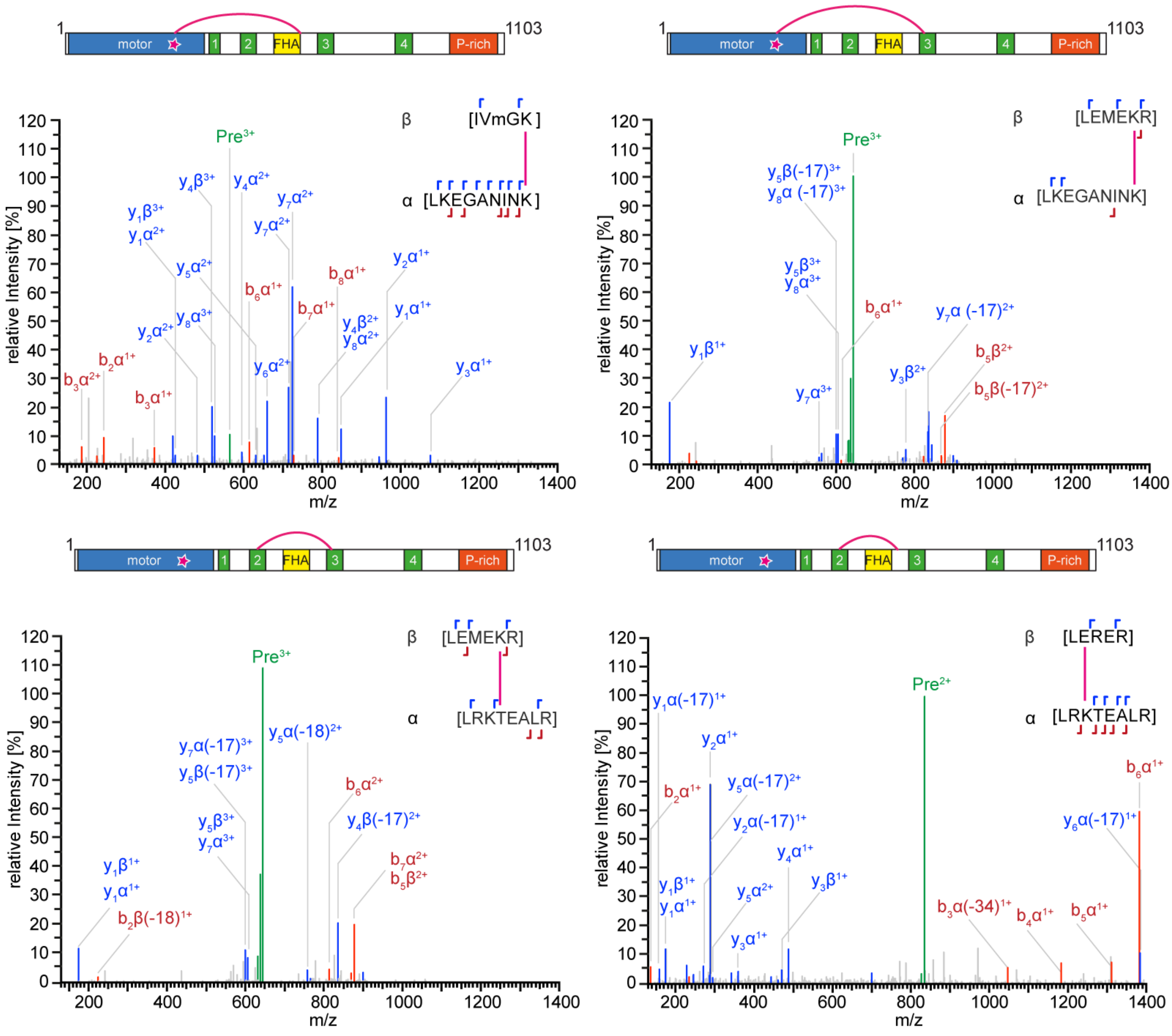


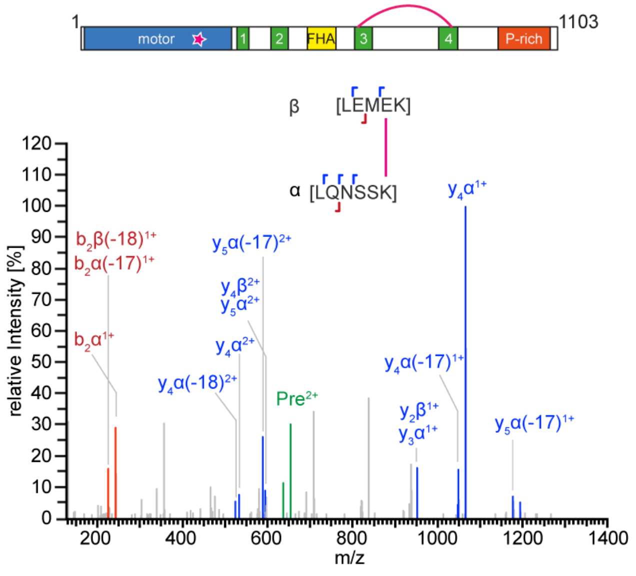


**Supplementary Figure 1: Verification of intramolecular crosslinks in KIF1C.**

Fragmentation spectra of crosslinked peptides as indicated in primary structure and peptide sequences above each plot. All ion fragments for which peaks could be allocated are indicated above and below the peptide sequence with red and blue hooks. The precursor ion is shown in green, the peaks are labelled in colour with alpha and beta b-ions labelled in red and alpha and beta y-ions labelled in blue. Major peaks are labelled with peptide fragment name.

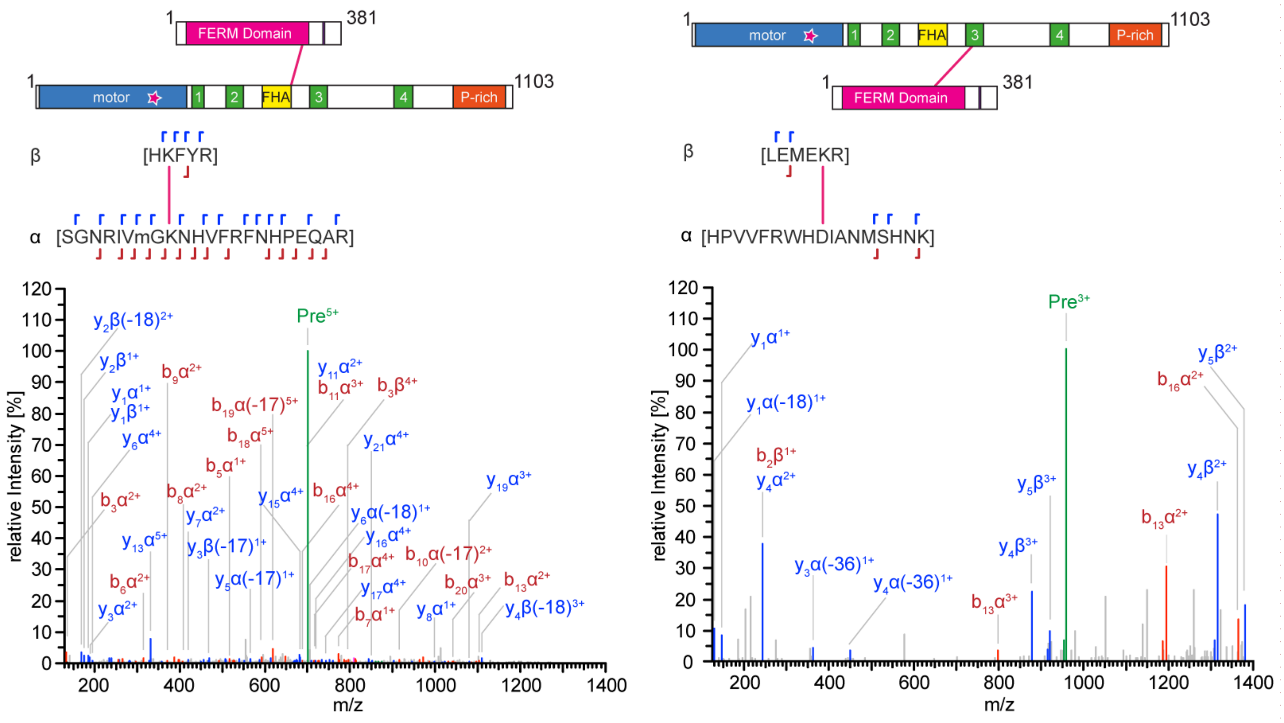


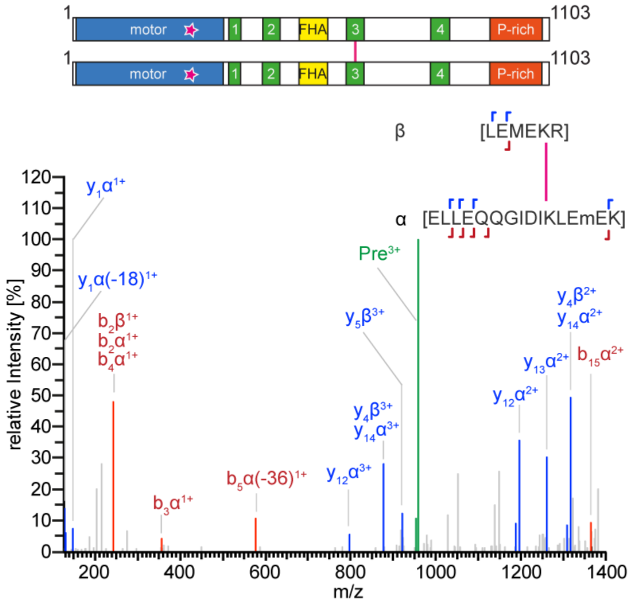


**Supplementary Figure 2: Verification of crosslinks between KIF1C and PTPN21 FERM domain.**
Fragmentation spectra of crosslinked peptides between KIF1C and PTPN21 adn in the KIF1C dimer as indicated in primary structure and peptide sequences above. All ion fragments for which peaks could be allocated are indicated above and below the peptide sequence with red and blue hooks. The precursor ion is shown in green, the peaks are labelled in colour with alpha and beta b-ions labelled in red and alpha and beta y-ions labelled in blue. Major peaks are labelled with peptide fragment name.
